# Insect-inspired, efficient event-based classification of tactile features

**DOI:** 10.64898/2026.06.18.733073

**Authors:** Lingsheng Meng, Kaushik Jayaram, Jean-Michel Mongeau

## Abstract

Tactile sensing enables humans and animals to detect and discriminate features during exploration and guide context appropriate actions. Compared to conventional touch sensors, sensing of tactile features in animals is fundamentally event-based through spikes. Yet how sensor mechanics shape spike activity for tactile perception is not well understood. Inspired by the American cockroach—an insect touch specialist—we developed a neuromechanical framework that linked antenna passive mechanics, mechanosensory encoding, and spike-based computation. A physics-based model of antenna bending simulated spatiotemporal strain patterns during contact, which were encoded into spike trains through a strain-to-firing mapping calibrated against electrophysiological recordings. The model captured antennal nerve activity observed *in vivo* by reproducing key features of population-level neural responses across multiple contact locations and speeds. Compared with conventional threshold-based encoding, the insect-inspired spike encoder preserved the spatiotemporal structure of tactile signals while achieving sparser activity. To establish a link between spiking activity and perception, we trained a spiking neural network to classify contact location and speed directly from the predicted spike trains. The network achieved >95% accuracy with reduced computational demands and enabled rapid discrimination within the first 170 ms of contact, indicating that sparse, event-based codes support fast and reliable tactile perception. Together, these results establish a mechanistic bridge between sensor mechanics and neural computation, revealing how physical interactions shape efficient sensory coding. This integrative framework advances our understanding of tactile perception and provides design principles for energy-efficient, neuromorphic tactile systems.

**Author Summary:** Animals use touch to explore their surroundings, identify objects, and make rapid decisions. Unlike most engineered touch sensors, which continuously transmit data, biological touch systems communicate through brief electrical signals called spikes. However, how the physical properties of a touch sensor influence these signals remains poorly understood. In this study, we used the antenna of the American cockroach as a model system to investigate how mechanics and neural activity work together during touch. We developed a computational framework that links the way an antenna bends during contact to the neural signals generated by touch-sensitive sensors. By comparing our model with neural recordings from living insects, we showed that it can reproduce key patterns of neural activity observed during tactile interactions. We found that the insect-inspired encoding strategy produces sparse signals that retain important information about where and how contact occurs. These signals enabled a neural network to rapidly and accurately identify contact location and speed while using fewer computational resources. Our results suggest that tactile perception emerges from a close interaction between sensor mechanics and neural processing. Beyond advancing our understanding of animal sensation, this work provides principles for designing energy-efficient touch sensors and neuromorphic robotic systems.

## Introduction

Insects exhibit remarkable efficiency, enabling them to navigate, explore, and interact with complex environments with little energy compared to current robots/AI. As such, insects have inspired more efficient AI for autonomous robots [1]. Among their sensory modalities, touch plays a crucial role in guiding rapid and slower exploratory behaviors such as tactile navigation [2–4], object localization and exploration [5–7], and rapid escape responses [8–11]. Antennae, the primary tactile organs of many insects, function as active sensors that probe the environment, extracting touch information through fine-scale interactions [4,6].

The performance of insect tactile systems arises from a tight integration of mechanical and neural processes, in which the geometry and material properties of the antenna shape the sensory input before neural encoding [12]. Recent studies have demonstrated that the mechanics of a sensor acts as a pre-neural filter that influences sensory processing, such as wing stiffness in detecting rotation during flapping [13] or antenna compliance in estimating obstacle distances and classifying objects during navigation [3,14,15]. Physical interaction dynamics between the antenna and its environment generate distributed strain fields that are transduced by mechanoreceptive sensilla into spatiotemporal spike trains. For instance, in the American cockroach *Periplaneta americana*, approximately 40,000 mechanoreceptive sensilla [16,17] distributed along the antenna transmit information via two primary antennal nerves [18] to the antennal mechanosensory and motor center (AMMC) [19,20]. Despite detailed anatomical and electrophysiological studies [14,18,21], the neural mechanisms which insects employ to efficiently encode and classify tactile features from high-dimensional strain inputs remain poorly understood. Additionally, robotic models of antenna have been created [22–24] with bioinspired mechanics to enhance tactile perception, with some approaching insect sizes [25,26]. These robotic sensors could benefit from bioinspired neural information processing to overcome the current limitations in computational efficiency for insect-scale platform integration.

Recent advances in neuromechanical modeling [27–29] and spiking neural networks (SNNs) [30] offer new opportunities to bridge this gap between insect sensor mechanics and neural computation. While many recent studies have provided important insights into sensory processing, a substantial portion of these works have focused on motor control and locomotion, where neural signals are often incorporated as predefined or task-specific inputs [31–34]. Furthermore, existing sensory models often rely on Spike-Triggered Average (STA) methods derived from white-noise mechanical inputs [35–37]. While these provide a linear approximation of sensory filters, the resulting rate-based waveforms lose spike timing information, which could be informative for sensory perception [38]. As a result, the mechanistic transformation from sensory input to neural encoding remains less explored in an integrated framework that explicitly links mechanics, receptor dynamics, and spike-based representations.

While SNNs are increasingly deployed in neuromorphic systems for tactile [39], visual [40], and olfactory [41] processing, their functional role in the integrated neuromechanics of sensory perception remains largely unexplored. Furthermore, it is not yet understood how insects utilize sparse, event-driven codes to achieve rapid tactile classification. Therefore, we hypothesize that antenna neuromechanics can generate sparse sensory codes for efficient tactile perception.

To evaluate how structural mechanics, neural encoding models, and network architecture each contribute to sparse sensory classification, in this work we developed a comprehensive neuromechanical framework that integrates physical simulation of antennal mechanics with a neural encoding model (**Fig 1**, **S1 Video**). Using a physics engine to simulate realistic antenna-object interactions [42], we converted distributed strain patterns into spike trains of campaniform and hair sensilla through a mapping calibrated against electrophysiological recordings. This approach allowed us to generate biologically plausible neural activity that preserves the spatiotemporal structure of tactile signals while achieving high sparsity. By training SNN classifiers directly on these predicted spike trains, we demonstrated that this integrated framework enables rapid discrimination of contact location and speed based on sparse, spike-based codes. Our results demonstrate that coupling realistic mechanics with neural computation enables efficient tactile feature discrimination, providing insights into the principles of energy-efficient processing in sensory systems and inspiration for bioinspired distributed tactile system design.

**Fig 1.**
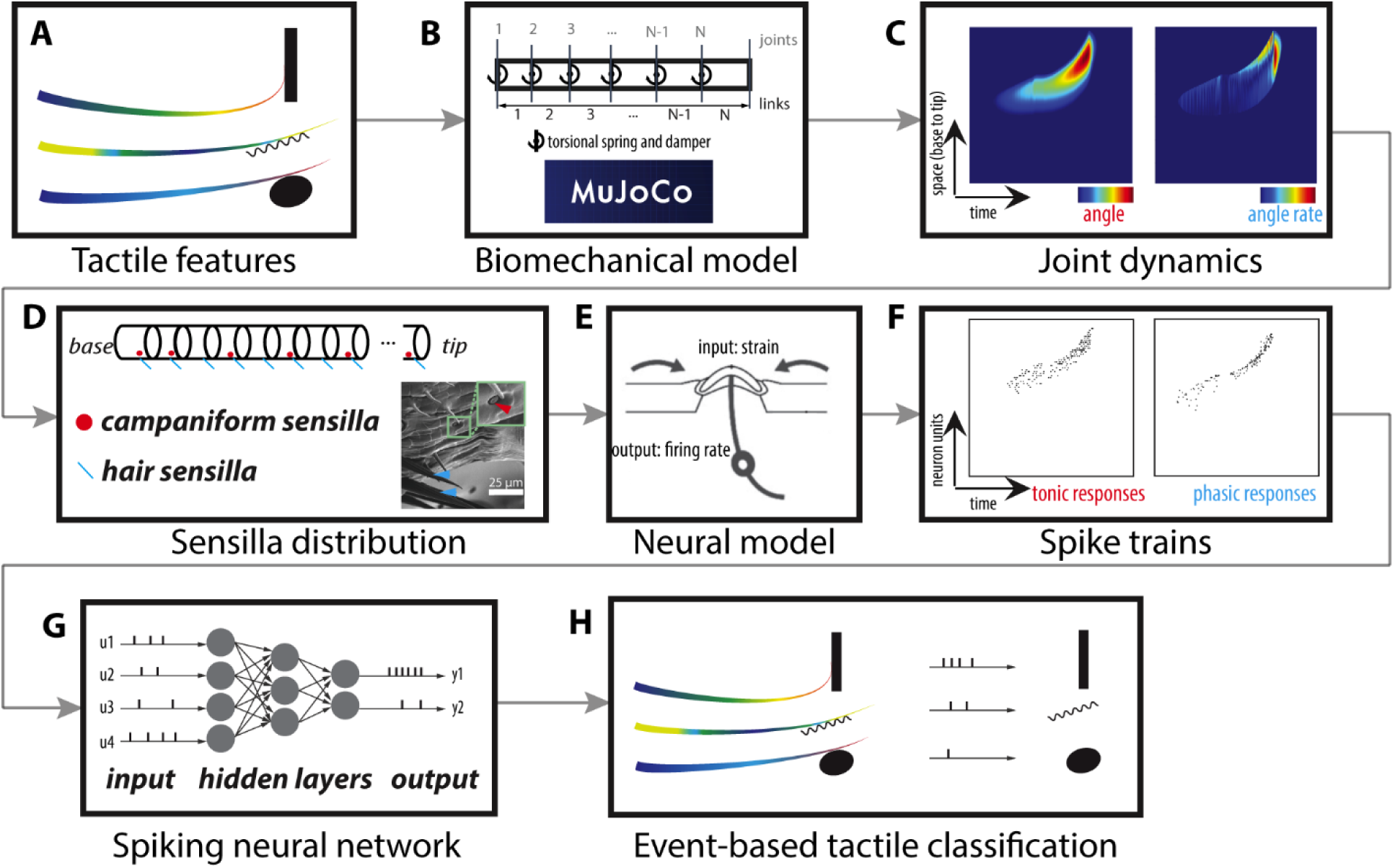
Development of a neuromechanical model of the antenna. (A) Example of tactile features. (B) Biomechanical model of the antenna represented as a kinematic chain, implemented in the MuJoCo physics engine. (C) Spatiotemporal strain patterns (tactile fields) generated during antennal deflections upon contact. Left: joint angles; right: joint angular velocities. (D) Distribution of mechanosensory sensilla along the antenna. (E) Phenomenological model of campaniform sensilla responses. (F) Predicted spike trains for campaniform (left) and hair (right) sensilla. (G) An example of SNNs. (H) Predictions from SNNs can be linked to distinct mechanical stimuli.

## Results

### Neuromechanical spike encoding preserves spatiotemporal tactile structure with sparse population activity

Antennal tactile encoding emerges from two key components: the mechanical filtering implemented by the structural and mechanical properties of the antenna, e.g. stiffness gradient, high damping, etc., and the neural filtering implemented by sensory receptors [12,21]. Thus, efficient tactile representation requires both appropriate sensor mechanics and biologically realistic neural encoding. Our previous work demonstrated that cockroach-inspired antennal mechanics enhance tactile discrimination by increasing the sparsity and spatiotemporal dispersion of strain fields across distinct mechanical inputs [43]. Building on this antenna modeling framework, we investigated whether a neuromechanical model grounded in cockroach antennal biomechanics could transform tactile interactions into spike-based population activity while preserving the spatiotemporal structure of strain dynamics (**Fig 2A, B**).

**Fig 2.**
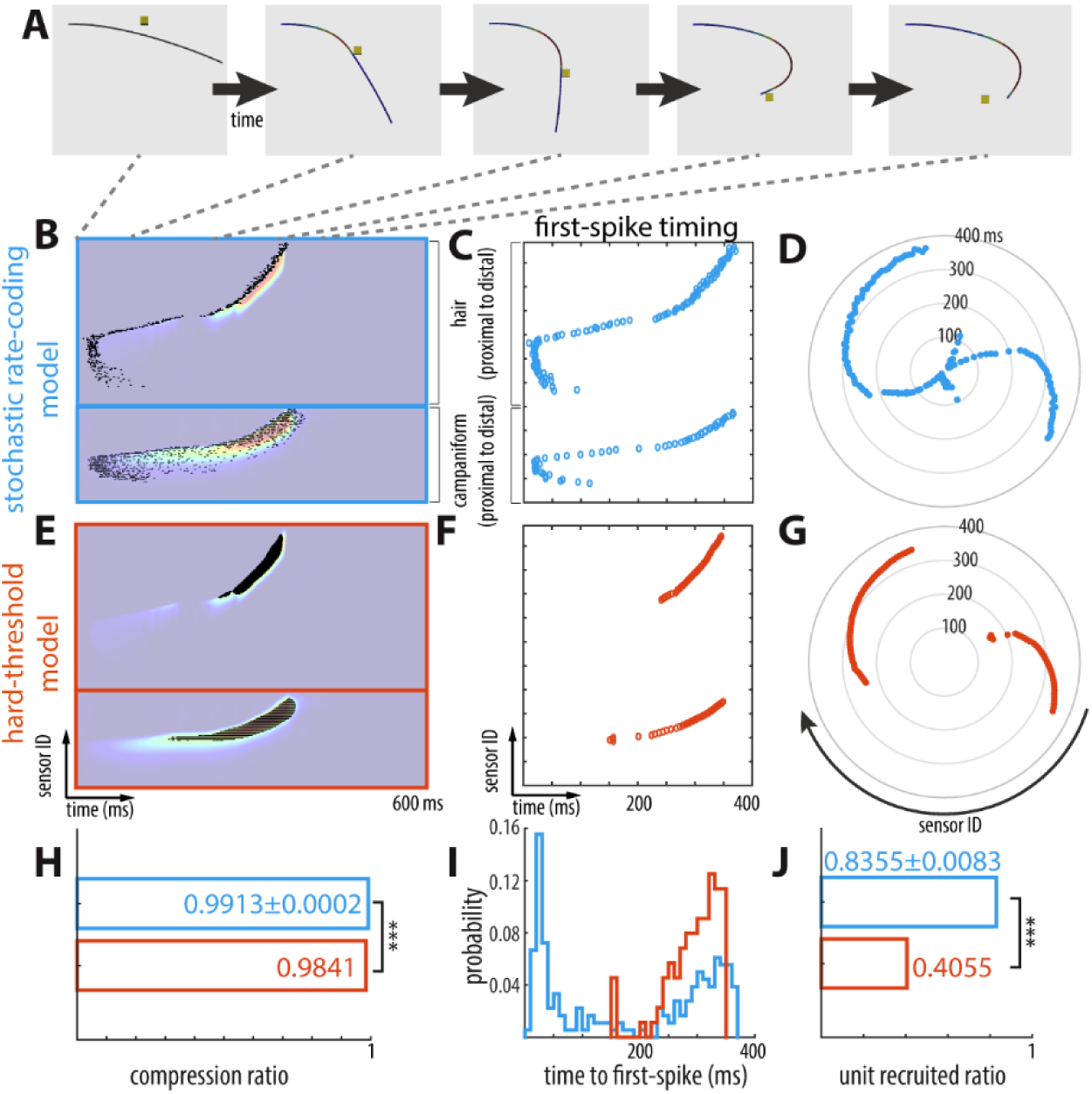
Neuromechanical spike encoding preserves tactile information with high efficiency. (A) Antenna dynamics in the *base-slow* condition. Frame sequences related to the tactile field in panel A and D. (B) Spike rasters generated by the stochastic rate-coding model overlaid on the tactile field (bottom: strain; top: strain derivative) generated by the biomechanical antenna model during the *base-slow* condition. (C) Time to first spike for all sensory units under stochastic rate-coding model. (D) Rose plot showing the distribution of first-spike timing across sensory units for the stochastic rate-coding model. (E-G) Corresponding spike rasters, first-spike timing, and rose plot for the hard-threshold encoding model (threshold = 0.5). (H) Compression ratio comparing stochastic rate-coding model (blue) and hard-threshold (orange) encoding. (I) Distribution of first-spike latencies for both encoding models. (J) Fraction of recruited sensory units under each encoding models. Statistical significance was determined via Wilcoxon signed rank test. ***: *p*<0.001. For panel H and J, standard deviation is shown numerically because errors bar would not be visible.

To evaluate neural encoding efficiency in the proposed antennal neuromechanical model, we compared spike trains generated by the insect-inspired stochastic rate-coding model with those produced by a conventional hard-threshold encoder applied to the same mechanically generated stain signals (**Fig 2E**). In a representative example (*base-slow* condition; see Methods), spike trains generated by the stochastic rate-coding model closely followed the spatial and temporal contours of the strain pattern (tactile field), capturing both the onset and offset boundaries of contact-induced strain and strain derivatives (**Fig 2B**). In contrast, the hard-threshold encoder (threshold = 0.5 × maximum amplitude) produced spikes only in regions of high signal magnitude, resulting in a loss of fine temporal structure and incomplete representation of the tactile field (**Fig 2E, G**).

Temporal response properties further distinguished the two encoding models. The stochastic rate-coding model exhibited shorter latencies to first spike across sensilla compared with threshold encoding (**Fig 2C, F, I**), indicating greater sensitivity to mechanical input and faster response initiation. Rapid spike generation is likely critical for insects, as is the importance of precise spike timing for mediating rapid escape responses [44,45]. Consistent with this notion, rose plots of first-spike timing revealed a faster, broader and more distributed temporal response in the stochastic rate-coding model (**Fig 2D, G**).

At the population level, the stochastic rate-coding model recruited a substantially larger fraction of responsive units than the threshold encoder across all 6 mechanical conditions studied in this work (*base-slow*, *base-fast*, *middle-slow*, *middle-fast*, *tip-slow*, and *tip-fast* in **S2 Video**) and for all tested threshold values (Wilcoxon signed rank test: *p*<0.001; **Fig 2J, S1 Fig**), indicating that more sensory channels contributed information about the stimulus. Despite this broader recruitment, the stochastic rate-coding model generated fewer spikes overall, resulting in a higher data compression ratio than the hard-threshold encoder (**Fig 2H, S1 Fig**). Sensitivity analysis of threshold values revealed that a lower threshold increases spike counts across a broader area of the tactile field (**S1 Fig**). Compared to the hard-threshold model, the stochastic rate-coding model exhibited a higher unit recruited ratio across all conditions for thresholds between 30% to 90%. Furthermore, the stochastic rate-coding model resulted in higher data compression ratio at thresholds of 30% to 60% and demonstrated faster response times in all conditions. Together, these results demonstrate that the stochastic rate-coding model achieves efficient tactile encoding, preserving essential stimulus features with sparser population activity.

### Electrophysiological recordings reveal diverse neural response patterns during antenna-object collisions

To validate predictions from the neuromechanical model, we performed extracellular recordings from the antennal nerve of *P. americana* during controlled antenna-object collisions (**Fig 3A**). After spike sorting (**Fig 3B**), we identified 39 units from 12 animals, and their Gaussian-convolved firing rates were extracted for further analysis (**Fig 3C**). Three distinct response types were observed across units (**Fig 3D**): (i) phasic onset responses activated at contact initiation, (ii) phasic offset responses occurring at the object detached from the antenna, and (iii) phasic-tonic responses sustained throughout the contact period. During the controlled collisions, the antenna accelerated with object upon contact onset, as expected from previous work [15], and reversed direction as the object detached (**S2 Video**), producing distinct mechanical phases. Units with purely phasic responses likely correspond to chemo-mechanosensory hair sensilla [21,46], which are velocity-sensitive and respond transiently to rapid deflections. The presence of both onset- and offset-tuned units may reflect the circumferential distribution of hairs around the antenna, such that different subsets are activated depending on the direction of bending. In contrast, phasic-tonic units, which remained active during sustained bending, are consistent with the activity profile of campaniform sensilla that encode strain in the antennal cuticle [21,47]. Analysis of response timing across units revealed variable delays to peak firing following contact onset (**S2-7 Figs**), suggesting spatially distributed sensory encoding of the antenna.

**Fig 3.**
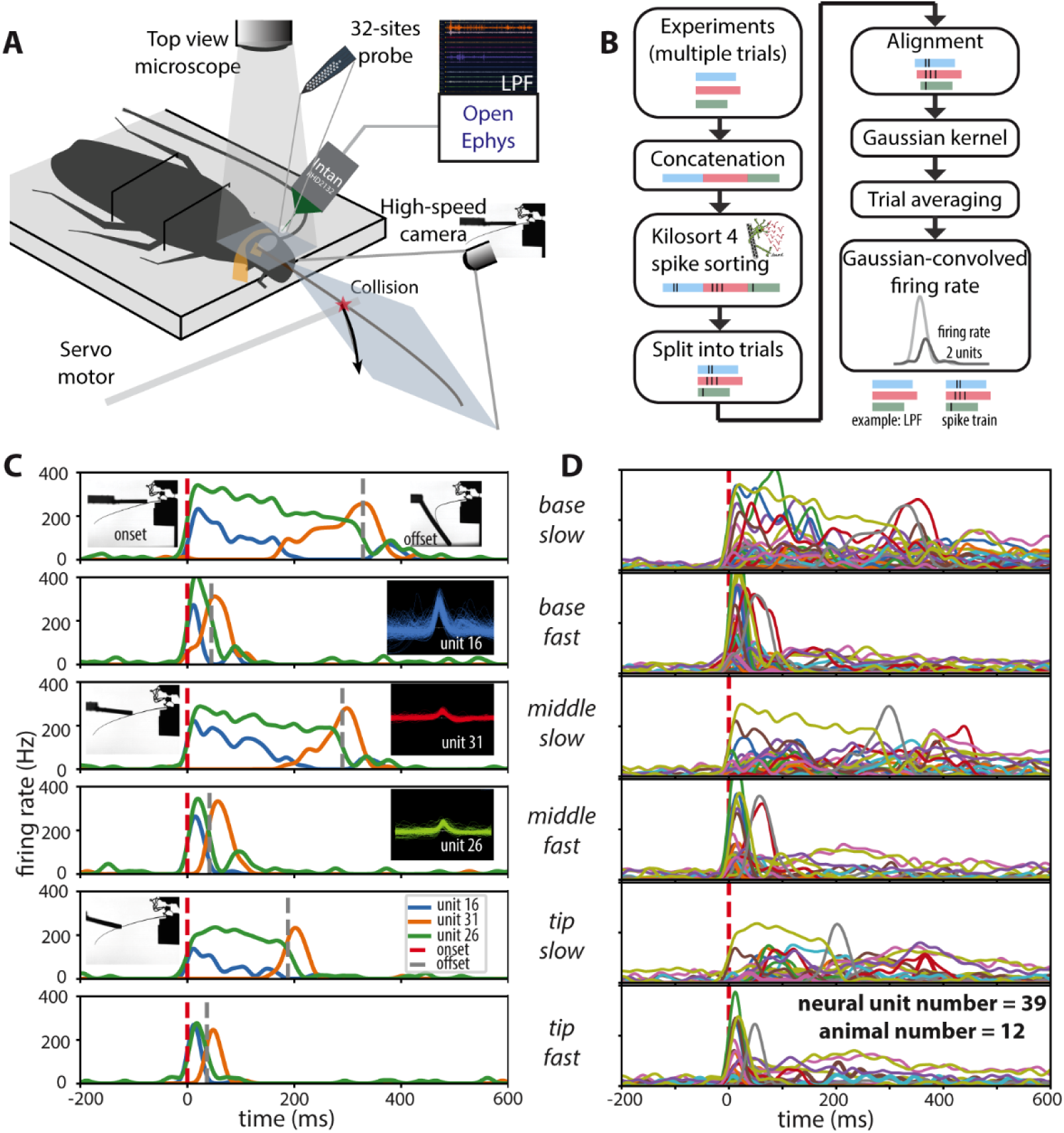
Extracellular recordings reveal diverse neural response patterns in the antennal nerve during controlled antenna–object collisions. (A) Experimental setup for electrophysiological recording. A probe was inserted into the exposed antennal nerve and connected to an Open Ephys acquisition system. A high-speed camera synchronized with the electrophysiology system recorded antenna–object collisions. (B) Post-processing pipeline for electrophysiological data. Recordings from multiple trials were concatenated and spike sorted. Sorted spikes were split into individual trials, aligned to contact onset, and smoothed using a Gaussian kernel to compute firing rates. (C) Population firing-rate responses across all recorded units (*n*=39 units from 12 animals) under six stimulation conditions. Each trace represents one sorted unit. The red dashed line indicates contact onset. (D) Examples of three distinct unit response types in the same recording under six stimulation conditions: *base-slow*, *base-fast*, *middle-slow*, *middle-fast*, *tip-slow*, and *tip-fast*. The red and gray dashed line indicates contact onset and offset, respectively.

### Population firing rate predicted by the neuromechanical model reflect electrophysiological recordings

To evaluate whether the neuromechanical model can reproduce neural population dynamics observed *in vivo*, we estimated population firing rates by combining the simulated responses of campaniform and hair sensilla (see Methods). The median firing rate across sorted units from electrophysiological recordings was used as an experimental reference to tune the neural model. The tuned parameters of the neuromechanical model exhibited clear biological interpretations (see Methods). The intercept parameter, which shifts the strain input to the neural model, determines whether the mechanosensory sensillae respond only after a strain threshold. A negative intercept indicates that strain must exceed a certain level before eliciting a neural response. The threshold parameter in the model represented the saturation level of the firing rate. During antenna-object collisions, each annulus typically bends from 1° to 10°, as measured from the tactile field, which is several orders of magnitude larger than typical tibial deflections from which the neural model was originally derived [48]. The tuned model yielded a relatively low threshold value, such that sensilla responses saturated at an annulus bending angle of 0.47°, well below the maximum observed deflections. This result suggests that mechanosensory sensilla operate near saturation during high-amplitude antennal movements, possibly to maintain sensitivity to strain direction rather than magnitude. The tuned ratio value between campaniform and hair sensilla contributions indicated a dominant role of hair sensilla in population-level encoding. This finding is consistent with the electrophysiological results, where less than 50% of recorded units exhibited phasic-tonic responses (**S2 Fig**). Given the higher density and broader distribution of mechanosensory hairs compared to campaniform sensilla —with a numerical ratio of approximately 32: 1 [16]—it is reasonable that hair sensilla contribute more strongly to the overall antennal neural output.

After parameter tuning, both the recordings and model predictions under slow contact speed conditions exhibited two distinct peaks corresponding to contact onset and offset (**Fig 4**). Overall, the tuned neuromechanical model successfully captured characteristic biphasic activity peaks and total response duration of the population-level neural responses (average MSE=0.004), supporting its validity as a framework for antennal mechanosensory coding.

**Fig 4.**
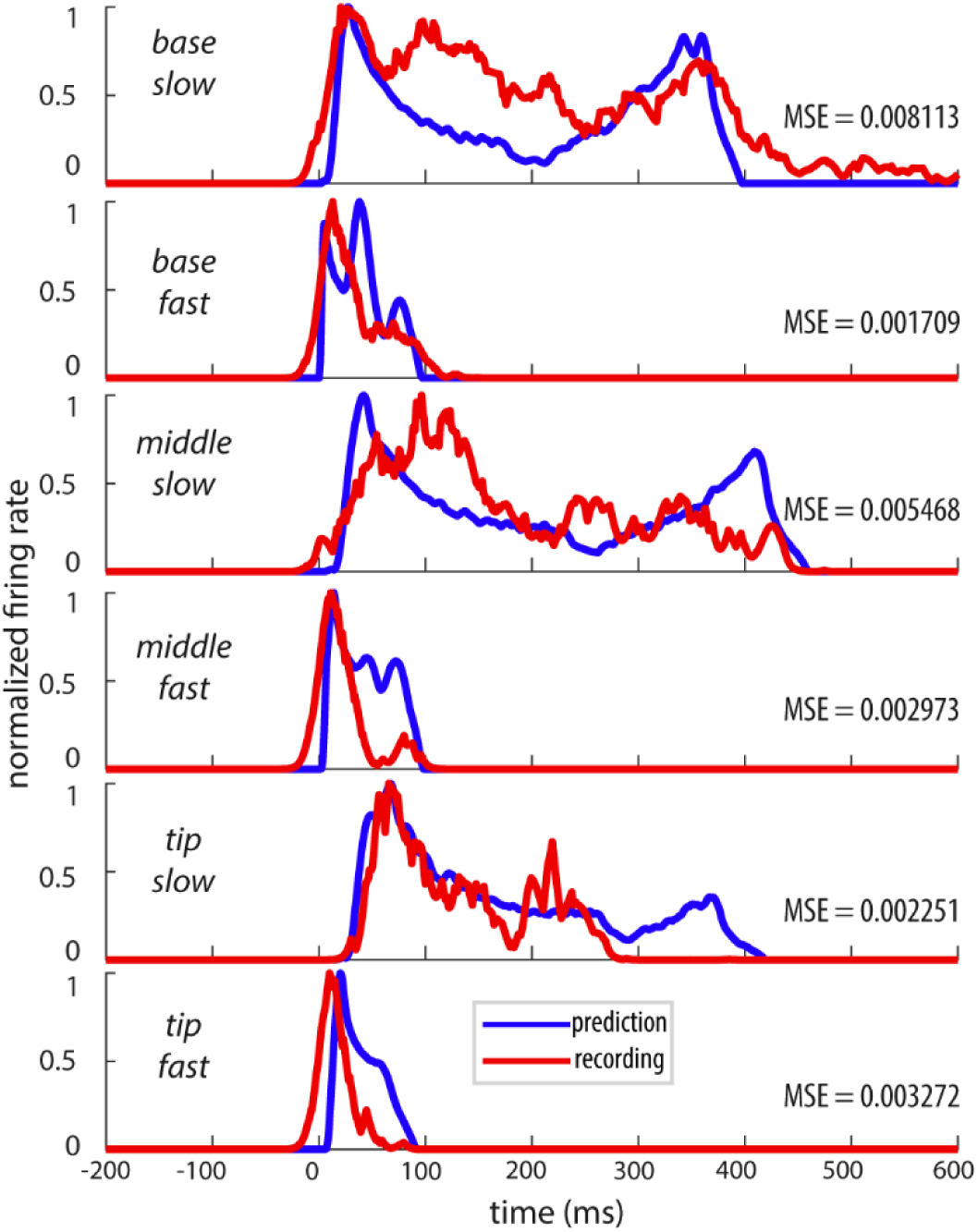
Comparison between predicted and recorded population firing rates during antenna-object collisions. Normalized population firing rates from the neuromechanical model (blue) after tuning and electrophysiological recordings (red) are shown for six stimulation conditions: *base-slow*, *base-fast*, *middle-slow*, *middle-fast*, *tip-slow*, and *tip-fast*. Each panel represents the average population response (N=39 units from recordings, N=217 units from the neuromechanical model) aligned to the onset of antenna–object contact (time = 0 ms).

### SNN discriminates antenna-object collision conditions from sparse neuromechanical codes

To evaluate whether the sparse, spike-based representations generated by the antennal neuromechanical model are sufficient for tactile discrimination, we trained a SNN directly on the predicted spike trains (**Fig 5A-D**; see Methods). The SNN framework was chosen for its biological plausibility and energy efficiency, as it processes information through discrete spike events rather than continuous activations [43]. Unlike conventional artificial neural networks that reply on dense Multiply-Accumulate Operations (MACs) at every timestep, spike-based networks operate in an event-driven manner, where synaptic updates are triggered only when spikes occur, thereby reducing effective computation under sparse firing conditions [49,50].

**Fig 5.**
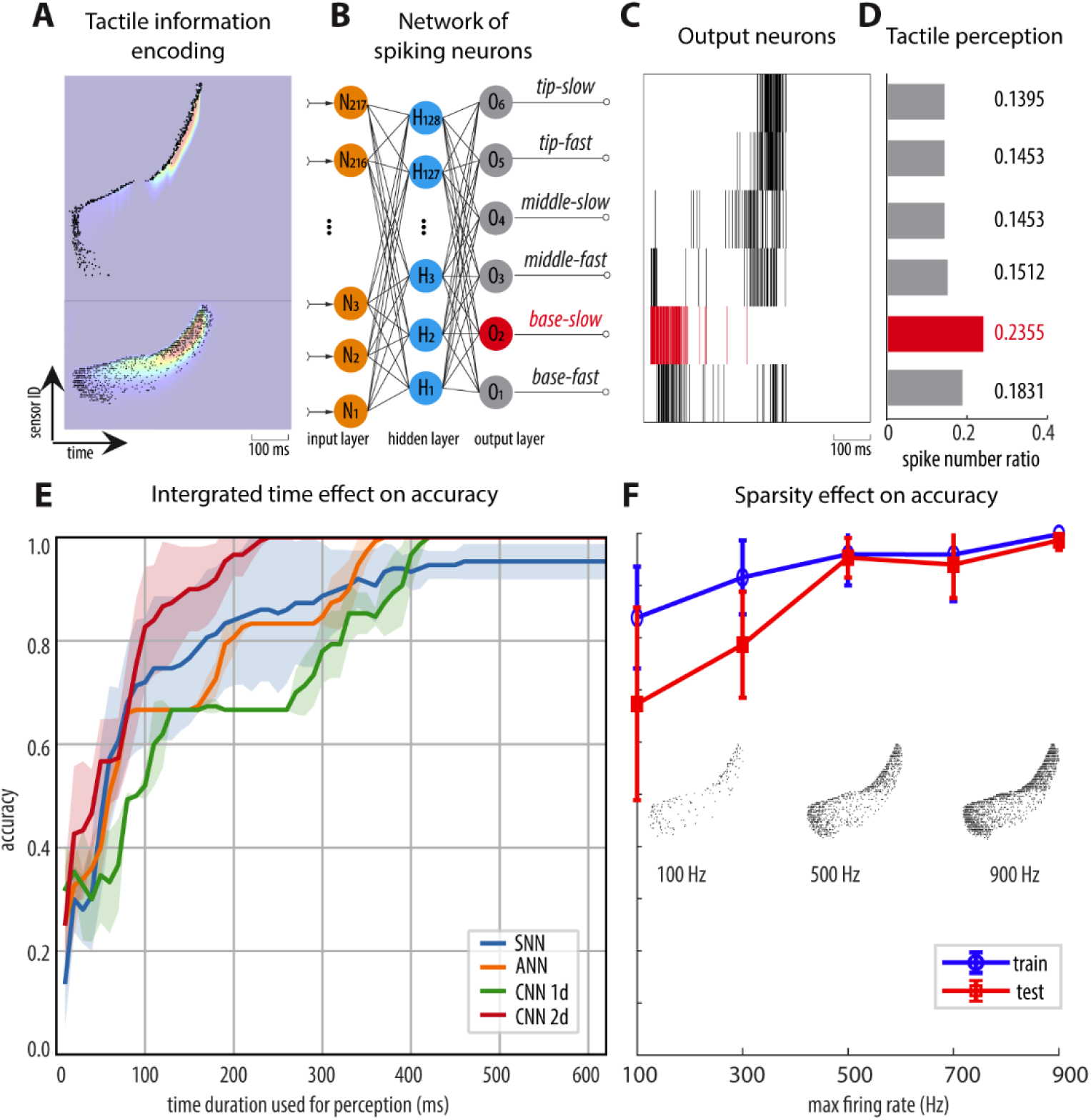
Spiking neural network decodes antenna–object collision conditions from sparse neuromechanical spike trains. (A) Example spike-based tactile representation generated by the neuromechanical model during *base-slow* condition. Spike rasters are overlaid on the corresponding strain (bottom) and strain-derivative (top) tactile fields. (B) Architecture of the SNN classifier. Spike trains were fed into a multilayer feedforward SNN composed of leaky integrate-and-fire neurons, with six output neurons corresponding to the six collision conditions. (C) Example output spike trains from the SNN during inference. (D) Classification was determined by the output neuron emitting the highest spike count over the integration window, illustrating correct classification of the base-slow condition. (E) Time-resolved classification performance. Shaded region indicates mean ± s.d. across five independent training runs. (F) Mean training and test accuracy (± s.d.) across datasets generated with varying maximum firing rates (100–900 Hz). Inset: representative spike patterns at each firing rate.

For each of the six antenna-object contact conditions (*base-slow*, *base-fast*, *middle-slow*, *middle-fast*, *tip-slow*, and *tip-fast*), 50 spike trains were generated using the stochastic spike generator with a maximum firing rate of 500 Hz (see Methods). The dataset was split into 90% for training and 10% for testing. Each model was trained independently five times to account for random initialization and stochasticity in optimization (**S8 Fig**). Across runs, the SNN converged rapidly within 40 training epochs, reaching an average training accuracy of 96.00 ± 5.97% and a mean test accuracy of 95.33 ± 3.80%. These results demonstrate that the neuromechanical spike encoding preserves sufficient spatiotemporal information to reliably discriminate tactile contact conditions.

Because SNN outputs are spike trains, classification was determined by identifying the output neuron with the highest spike count over the evaluation window (**Fig 5C, D**). Consequently, classification performance depends on the temporal integration window. To evaluate the system’s capacity for rapid and dynamic tactile inference, we computed time-resolved classification accuracy by progressively increasing the integration window from contact onset to the full duration of the tactile tensor. Classification accuracy increased with longer integration times, reflecting the accumulation of additional sensory evidence, while training itself was optimized using the full temporal window (**Fig 5E**). Notably, the SNN achieved 80% classification accuracy within the first 170 ms following contact which is corresponding to only ∼30% of the total tactile field, highlighting the system’s ability to rapidly infer contact conditions from early sensory spikes. To benchmark performance against alternative classifiers, we trained a 1D convolutional neural network (CNN), a 2D CNN, and a conventional artificial neural network (ANN) using the same spike trains generated by the neuromechanical model (**Fig 5E**, **Table 1**). Because these classifiers require continuous input, spike trains were converted into 32-bit floating-point representations containing binary values (0 and 1). The static classifiers (ANN and CNNs) achieved a peak accuracy of 100.0%, whereas the SNN reached a test accuracy of 95.3%. However, the SNN exhibited superior performance under short temporal integration windows, achieving reliable early-time classification with substantially lower model complexity. To quantify computational efficiency, we estimated Multiply-Accumulate operations (MACs) as a proxy for energy consumption. The ANN operated on time-averaged spike inputs, requiring a single forward pass per sample. In contrast, the SNN processed spikes sequentially across all time-steps. Despite this temporal iteration, the event-driven nature of spike computation of the SNN significantly reduced effective MACs because synaptic updates occurred only when spikes were present. Under the measured spike sparsity, the effective computational cost of the SNN was approximately five times higher than that of the ANN and remained substantially smaller than that of the CNN architectures. Together, these results highlight the computational efficiency and temporal sensitivity of spike-based processing, demonstrating that sparse neuromechanical codes can support accurate and rapid tactile discrimination with reduced computational demand.

**Table 1.** Comparison of classifiers for antenna-object collision discrimination.

| Classifier | SNN | ANN | 1D CNN | 2D CNN |
| --- | --- | --- | --- | --- |
| Input type | Spike trains | Float 32 | Float 32 | Float 32 |
| Overall accuracy (%) | 95.3 | 100.0 | 100.0 | 100.0 |
| 80% accuracy time<br>(ms) | 170 | 190 | 320 | 90 |
| Model Size<br>(Parameters) | 28.7 K | 28.7 K | 41.5 K | 8.65 M |
| Model memory<br>(KB) | 342 | 342 | 494 | 101389 |
| MACs | 140.6 K | 28.5 K | 0.6 M | 13.0 M |
| Inference type | **Event-driven** | Static | Static | Static |

Finally, we examined how neural sparsity influences classification performance by systematically varying the maximum firing rate of the spike generator to produce datasets with different spike densities (**Fig 5F**). When the spike trains were extremely sparse (100 Hz maximum rate), classification accuracy was lowest. As firing rate increased, accuracy improved, suggesting that denser spike activity provides richer temporal and spatial information for discrimination. However, the accuracy plateaued at ∼500 Hz, suggesting a diminishing return on classification accuracy with higher spike density.

## Discussion

In this study, we developed and validated a neuromechanical modeling framework that links antennal mechanics with neural population activity in insect. By integrating a biomechanical model of antennal bending with a phenomenological neural encoding model of insect mechanoreceptors, we predicted the population firing rate responses during antenna-object collisions. The predicted responses captured the dynamics of electrophysiological recordings by capturing key temporal features such as phasic onset and offset peaks. Model tuning revealed biologically interpretable parameters that reflect sensillum threshold, sensitivity, and relative contributions of different receptor types. Furthermore, spike trains generated from the neuromechanical model were used to train a SNN for tactile classification, demonstrating that sparse, biologically grounded sensory codes can effectively support event-based tactile discrimination tasks. Together, these results provide a quantitative bridge between mechanical interactions, neural encoding, and downstream computational processing, offering new insights into the mechanistic basis of tactile sensing in insects and the development of bioinspired tactile sensing systems.

### Mechanocomputation and physical intelligence

Our results demonstrate that the cockroach antenna functions as a physically intelligent system, where sensory processing is embedded directly within its mechanical structure. Rather than passively transmitting forces to neural receptors, the antenna’s exponential stiffness gradient acts as a mechanical pre-filter that transforms raw tactile interactions into spatially structured patterns before neural transduction [43]. This finding aligns with the concept of morphological computation, in which body mechanics offload computational demands [51]. The exponential decreasing stiffness profile of the antenna [15] enhances sparsity and separability of strain fields across contact conditions, effectively preprocessing tactile information at the physical level. In contrast, alternative stiffness profiles produced strain patterns with reduced dynamics range [43]. Thus, morphology directly constrains the operating regime of the sensory network.

### Mechanosensory contributions to antennal tactile encoding

The insect antenna is a versatile sensory organ that integrates input from multiple types of mechanosensory sensilla to support tactile exploration and perception [4]. In this study, we modeled only two primary classes of sensilla—mechanosensory hairs and campaniform sensilla —as the main contributors to antennal tactile encoding within the flagellum. However, other mechanoreceptors distributed along the antenna base (scape and pedicel) may contribute to the overall encoding of antenna–object contact. In particular, hair plates located near the base of the antenna provide proprioceptive feedback related to joint angle and movement direction, which could play a role in encoding the position and movement direction of the antenna during tactile sampling [5,52]. Similarly, chordotonal organs embedded within antennal segments can detect stretching and vibration, providing dynamic information about antennal movement and mechanical resonance [53,54]. These sensors could influence tactile encoding through high-frequency mechanical oscillations that arise during vibration transmission of antenna-object contact, although these oscillations are likely small due to the high damping of the antenna [15]. Neural responses to mechanical waves propagating through the antenna shaft may therefore contribute to population activity patterns recorded but were not represented in the present model. Integrating these additional mechanosensory modalities into future models could provide a more complete understanding of mechanical encoding in the antenna.

### Limitations of the current neuromechanical model

Several assumptions made in the present model limit its biological realism. Structurally, the biomechanical model represented the antenna as a two-dimensional serial linkage composed of rigid annuli connected by compliant joints. This simplification ignores out-of-plane bending and the complex anisotropic properties of the cuticle along the annulus length [14]. For computational simplicity, each annulus was assigned only one representative hair sensillum and one campaniform sensillum, although anatomically, dozens of mechanosensitive hairs and roughly three campaniform sensilla occur non-uniformly per annulus [16–18,55]. This simplification likely diminishes the realism of spatial encoding and may lead to an underestimation of population-level response diversity.

On the neural modeling side, the phenomenological equations used to describe campaniform and hair sensillum dynamics were originally derived from tibial mechanoreceptors [48]. Although this approach captures key phasic-tonic features, it does not fully replicate their biological responses. For example, the model predicts vigorous discharges following force decrements while actual campaniform sensillum responses are smaller in magnitude and often delayed. Furthermore, regional variations in campaniform sensillum properties likely influence model performance. Specifically, variations in sensillum morphology [56,57] and cuticular material properties [58,59] collectively modulate mechanical transduction and thus limit the model’s generalizability. In addition, due to the absence of a dedicated neural model for mechanosensitive hairs, the campaniform sensillum model was adapted to approximate hairs’ behavior. Developing a more biologically accurate neural representation of hair sensilla could improve the fidelity and predictive power of the neuromechanical model.

### Discrepancies between model predictions and electrophysiological recordings

While the neuromechanical model reproduced many key features of antennal population responses, discrepancies remained, especially near the stimulus offset. Neural activity decayed more rapidly to baseline, while the model exhibited a slower return (**Fig 4**). Several factors could account for this difference. Biologically, the current model includes only campaniform and hair sensilla and assumes simplified receptor distributions (see previous section), thereby excluding other mechanosensory organs that may contribute to faster transient responses *in vivo*. Additionally, the simplified strain-to-firing mapping in the model is governed by a saturating linear mapping followed by a stochastic spike generator. This formulation lacks spike-frequency adaptation and therefore may overemphasize tonic encoding components, leading to delayed or prolonged activation following contact (**Fig 4**). Experimental limitations may also contribute to these discrepancies. Extracellular recordings of antennal nerve represent compound population signals that integrate activity across multiple receptor types and may include unmodeled sensory units like chordotonal organs in this study. Such recordings may emphasize rapidly adapting components or underrepresent slowly decaying activity from campaniform sensilla. Anatomically, the cockroach antenna contains two parallel afferent tracts—anterior nerve and posterior antennal nerves—which merge within the scape [55]. These tracts carry spatially segregated afferents from the anterior and posterior surfaces of the flagellum [60]. In this study, the recording electrode was positioned dorsally, likely overrepresenting signals from specific antennal mechanoreceptor populations.

### Sparse and spatiotemporal coding as a general principle of tactile sensing

The neuromechanical model developed in this study provides insight into how insect antennae transform mechanical interactions into sparse and spatiotemporally structured neural representations. In this framework, distributed strain patterns are firstly generated by the biomechanical model based on antennal mechanics. These mechanically filtered signals are then converted by the neural encoding model into temporally precise spike trains across mechanosensory units, resulting in a population code in which only a subset of neurons is active at any given moment (**Fig 2B**). Such sparse coding is widely recognized as a fundamental principle in sensory systems. By restricting activity to a limited number of neurons, sparse representations increase coding capacity, facilitate efficient downstream decoding, and minimize metabolic cost [61].

In the antennal system, such sparsity could enable rapid detection of contact events and efficient discrimination between distinct tactile features without requiring continuous high-frequency firing. This efficient encoding is particularly advantageous for learning in insects, where sensory processing operates under severe energy and computational constraints [1,62]. The phasic and phasic–tonic response patterns observed in both our electrophysiological recordings and model predictions exemplify this efficiency: phasic units provide temporally precise markers of contact onset and offset, whereas tonic units encode sustained deformation and object interaction dynamics. Together, these features could form a compact yet high-fidelity neural representation for tactile exploration.

Similar spatiotemporally structured coding strategies are well documented in the vibrissal (whisker) system of rodents, one of the most studied models of active touch. In rats, deflection of whiskers elicits precise and millisecond-scale responses in primary afferents and neurons of the somatosensory barrel cortex which are tuned to the position, velocity and direction of whisker motion [63–65]

Selectivity for tactile features is maintained along the entire sensory pathway, from the brain stem to the somatosensory barrel cortex, allowing object location and texture to be encoded with high temporal precision [66,67] Similar to the insect antenna, the whisker array functions as a biomechanical filter that converts external forces into strain patterns on the whisker follicles. This suggests that neuromechanical coupling may represent a general mechanism for tactile sensing across species [68,69].

Analogous principles are also observed in the human tactile system, where mechanoreceptors in the glabrous skin exhibit sparse, highly informative firing patterns during contact events. Rapidly adapting (RA) and Pacinian (PC) afferents respond phasically to transient contact and vibration, encoding dynamic changes in indentation, whereas slowly adapting (SA) afferents sustain firing during static pressure to represent spatial features such as shape and texture [70]. These encoding properties collectively support energy-efficient and temporally precise perception of texture, shape, and motion [29,71].

### Performance and future directions of SNN decoding

Our SNN classifier demonstrated that spike-based representations generated from the neuromechanical model are sufficient for discriminating different antenna–object collision conditions, including object location and speed. The high classification accuracy achieved with sparse input supports the idea that biologically inspired sensory codes can efficiently drive event-based neural computation. Nonetheless, the current SNN architecture represents a relatively simple feedforward topology. Future work could explore more advanced spiking architectures, such as deeper layers, recurrent spike neural networks [72], and modulated spike-timing-dependent plasticity-based learning [73], which may better capture temporal dependencies and adaptive tactile patterns. Additionally, training protocols could be enhanced by introducing temporal jittered inputs, variable contact durations, and noise-perturbed stimuli to improve robustness and generalization to previously unseen contact scenarios. These specific augmentations can mimic the variability of natural tactile interactions and align with biological neural adaptation, in which neural circuits continuously adapt to variable tactile experiences.

### Neuromorphic implementation and broader applications

The integration of a neuromechanical model with SNN decoding provides a framework readily transferable to neuromorphic hardware for real-time tactile processing. Implementing such systems on neuromorphic chips would enable energy-efficient, event-driven computation directly at the tactile sensor system, reducing communication latency and power consumption enabling integration onto insect-scale platforms [74]. This approach could inform the design of bio-inspired event-based tactile sensors, such as artificial antennae or hair-like sensory arrays for autonomous robots [25]. These systems could perform local contact detection and surface texture discrimination using sparse spiking outputs, analogous to insect mechanosensory codes. Ultimately, the neuromechanical integration demonstrated here not only proposes an architecture for insect tactile perception but also lays a foundation for next-generation tactile sensing and control systems in robotics and neural engineering.

## Methods and Materials

### Biomechanical model of antenna for the transmission of tactile stimuli

Inspired by the morphology and mechanics of the antenna flagellum of the American cockroach *P. americana* [14,15], we developed a biomechanical model comprising 140 serially connected rigid links linked by hinge joints (**Fig 1B**). The stiffness *k* and damping *c* of each joint *n* were defined as exponential functions to capture the mechanical stiffness gradient from the base to the tip of the flagellum [15].

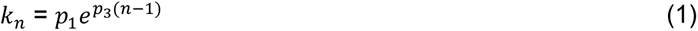

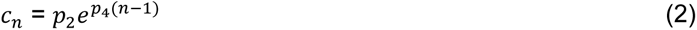

where *p*_1_ and *p*_2_ are the stiffness and damping constants at the basal joint, respectively. The parameters *p*_3_ and *p*_4_ define the exponential decay rates of stiffness and damping along the flagellum. Detailed derivation and validation of the parameterization can be found in previous work [43]. The antenna model was implemented using the MuJuCo physics engine [42], which provides a computationally efficient and accurate framework for simulating multi-body dynamics in contact-rich environments.

### Generation of spatiotemporal strain fields from the biomechanical model

Joint angle trajectories of the biomechanical antenna model were recorded within the MuJoCo physics engine as the simulated antenna contacted tactile features. Assuming a linear relationship between joint rotation and the local strain transmitted to mechanosensory sensilla [14], each joint represented a spatially distributed strain sensor. Consequently, the arrangement of joints encoded spatial information, while sequential time steps captured the temporal dynamics of the strain response. Together, these components formed distinct spatiotemporal strain fields corresponding to different tactile interactions.

To reproduce conditions comparable to the electrophysiological experiments, the initial curvature of the virtual antenna was adjusted to match the natural resting curvature observed in *P. americana* (**S9 Fig**). A contact object identical in size to that used in the electrophysiology setup was simulated to collide with the antenna at a similar location and velocity.

### Mechanosensory sensilla distribution and neural modeling

Two types of mechanosensory sensilla located on the antennal flagellum were incorporated into the model: hair sensilla and campaniform sensilla. Mechanosensory hair sensilla exhibit phasic responses to mechanical stimulation and are primarily sensitive to contact velocity [21,46]. In contrast, campaniform sensilla, embedded within the cuticle at the junctions between annuli, display phasic-tonic responses and function as strain gauges that encode cuticular deformation [47,75].

Adult *P. americana* possess approximately 6,500 mechanosensory hairs, arranged in circumferential rings on each annulus [16,17]. For computational simplicity, the neural model included one representative hair sensillum per annulus, consistent with the spatial resolution of the biomechanical model at the annular level. Campaniform sensilla are distributed more sparsely: they occur on every annulus from the base through around 15^th^ annulus, and on every other annulus beyond that [18,76]. Accordingly, one campaniform sensillum was modeled per annulus for the first 15 annuli and on alternating annuli thereafter. To simulate neural encoding, we employed a phenomenological model originally developed for campaniform sensilla on the cockroach tibia [48]. This nonlinear model describes the transduction of mechanical input *u*(*t*) into a neural output *y*(*t*) using a state-space formulation derived from dynamic systems theory:

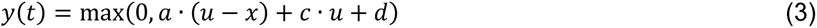

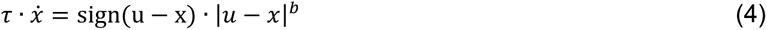

where *a, b, c, d* and *τ* are constant parameters. The interaction between the input *u* and the internal state variable *x* determines the phasic dynamics, while the tonic component is captured by the steady-state term *c* · *u* + *d*. Equilibrium occurs when *u* = *x*, at which point only the tonic term contributes to the output. Model parameters were tuned to reproduce the ramp-and-hold response characteristics of cockroach tibial campaniform sensilla [48]. To represent hair sensilla, parameters *c* and *d* were set to zero, thereby eliminating the tonic response and preserving purely phasic behavior. Finally, population neural activity was obtained by averaging the predicted firing rates of all modeled sensilla, incorporating axon conduction delays proportional to the distance between each sensillum and the antennal base, assuming an average conduction velocity of 2 m s^-1^ [77,78].

### Neuromechanical model tuning

Because the output of the biomechanical antenna model represents the joint angles of each annulus (in radians), whereas the input to the phenomenological neural model of campaniform sensilla corresponds to externally applied force on the cockroach tibia (in milliNewtons) [48], these two variables differ in scale and physical domain. Campaniform sensillae function as biological strain detectors, responding to local cuticular deformation that is linearly related to the external load on the cuticle [47,75]. Based on elastic bending analyses of insect legs, the relationship between the applied force and the resulting strain at the sensillum location can be approximated as linear [79]. Similarly, finite-element simulations of antennal bending demonstrated that local strain near the sensilla varies linearly with the joint angle [14]. However, the strain magnitude estimated from the antenna was approximately two orders of magnitude greater than that calculated in tibial models, precluding a direct transfer between systems. Therefore, in this study, we assumed a linear mapping between the joint angle output from the biomechanical model and the effective force input to the neural encoding model, while preserving the original internal parameters of the campaniform sensillum neural model. This mapping was expressed as:

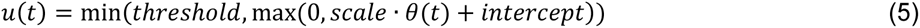

where *u*(*t*) denotes the equivalent mechanical input to the neural model, *θ*(*t*) is the joint angle, and scale, intercept, and threshold are scaling parameters determined to ensure data range consistency between model domains.

The scale parameter converts angular displacement (in radians) into an equivalent force range (in milliNewtons); for example, a 5° deflection (0.087 rad) scaled by 100 corresponds to approximately 8.7 mN. The intercept parameter (ranging from −0.4 to 0 mN) introduces a baseline offset to account for the activation threshold of the sensillum. This range was chosen to ensure the sensillum remains quiescent during minimal deflections (0 to 0.23°), preventing the model from generating spurious spikes at rest. To preserve the original dynamics of the campaniform sensillum neural model—since this study focused on validating the neuromechanical framework rather than modifying the underlying neural models—the threshold parameter was constrained between 0.3 and 4 mN [48], representing the maximum strain that the sensilla could effectively encode when the sensilla reach saturation frequency.

To tune model parameters and ensure that the predicted population firing rates from the neuromechanical model captured the dynamics of electrophysiological recordings, we minimized the mean MSE between the model prediction and experimental data across six mechanical conditions: *base-slow*, *base-fast*, *middle-slow*, *middle-fast*, *tip-slow*, and *tip-fast* (**S2 Video**). The tuning was performed using the Standard Particle Swarm Optimization algorithm implemented in MATLAB. It was selected over gradient-based methods because of its ability to navigate the non-convex, non-differentiable cost landscapes. The tuning was configured with a swarm size of 50 and a limit of 30 iterations, with a stall tolerance set to 5 iterations to ensure convergence. A fixed scaling factor (scale) of 100 was applied to establish a consistent baseline mapping between angular displacement and the input range of the neural model, while three free parameters were tuned: (1) the intercept and threshold used in the translation from joint angle to neural input, and (2) the ratio between campaniform sensilla (*f_c_*) and hair sensilla (*f*_ℎ_) contributions to the total population firing activity (*f_p_*). The combined population firing rate was computed as:

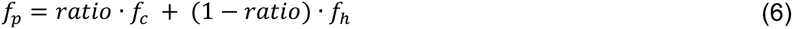

where *f_p_* denotes the predicted population firing rate, *f_c_* the normalized activity of campaniform sensilla, and *f*_ℎ_ the normalized activity of hair sensilla.

By minimizing the average mean squared error (MSE) between the simulations and experiments for all six conditions: *base-slow*, *base-fast*, *middle-slow*, *middle-fast*, *tip-slow*, and *tip-fast*, we tuned the intercept (-0.250) and threshold (0.575) used in the translation from joint angle to neural input, and the ratio (0.424) between campaniform sensilla and hair sensilla contributions to the total population firing activity.

### Spike generation from the neuromechanical model

To transform the continuous predicted neural activity into discrete spike representations suitable for SNN classification, we converted normalized firing rate into binary spike trains using a stochastic sampling framework. Each dataset produced by the neuromechanical model comprised a normalized firing-rate matrix (*T* × *N*), where *T* is the number of time steps and *N* is the number of modeled sensilla neurons. Spike generation was implemented through a Bernoulli sampling process, where the probability of emitting a spike at each time step was proportional to the instantaneous firing rate. To improve generalization and mimic biological variability, five forms of stochastic augmentation were applied:

1. Neuron-wise gain variability: To mimic heterogeneous neuronal sensitivity, a random multiplicative gain was assigned to each neuron, sampled from a log-normal distribution (σ=0.05) [80]. This modulation altered the amplitude of firing-rate responses while preserving the overall structure.
2. Temporal latency shift: Each sensillum’s spike train was randomly shifted in a uniform distribution U(-4, 4) ms to simulate global latency variations in sensory transmission due to various antenna length.
3. Spike probability sampling: The normalized firing rate was scaled by a maximum firing frequency (500 Hz unless otherwise specified, which was chosen to reflect the absolute refractory period of 2 ms in insect afferents [81,82] and multiplied by the bin width (1 ms) to obtain the spike probability per bin. Spikes were independently generated from a Bernoulli process.
4. Spike dropout: To model spontaneous transmission loss, a small fraction of spikes (2%) was randomly removed from each spike train.
5. Spike timing jitter: To simulate neural noise, a subset of spikes (5%) was randomly selected and shifted by ±2 ms.

Each augmented dataset generated a binary spike tensor (*T* × *N* × *K*), where *K* is the number of repeated realizations per stimulation condition (**S10 Fig**). This process provided stochastic yet physiologically realistic spike patterns, enabling the SNN classifier to learn temporally robust representations of tactile stimuli while maintaining the underlying structure of sensory coding.

### Compression ratio of spike-based encoding

To quantify the efficiency of spike-based tactile encoding, we computed a compression ratio (CR) that measures the reduction in memory required to represent tactile signals using sparse spike events relative to storing the full continuous-valued sensory data. The raw tactile field consists of continuous-valued strain and strain-derivative signals sampled at fixed temporal resolution across all sensory units. Storing such data requires allocating memory for every sensor at every time step. In contrast, spike-based encoding represents neural activity as discrete events in time, where information is stored only by the occurrence of spikes. This spike-based representation is inherently sparse. Thus, the compression ratio provides a principled metric to evaluate how efficiently the spike-based representation preserves information while reducing storage requirements. It was defined as the fractional reduction in memory usage achieved by spike-based encoding relative to the raw continuous representation:

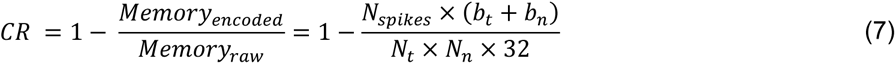

where the raw tactile data consist of *N_n_* sensory channels sampled over *N_t_* time steps. Each sample is stored as a 32-bit floating-point value. In the spike-based encoding, neural activity is stored as a list of spike events. Each spike is represented by a time index, requiring *b_t_* = 16 bits and a neuron index, requiring *b_n_* = 8 bits. *N_spikes_* is the total number of spikes in a trail. A higher compression ratio indicates a more memory-efficient encoding and sparser neural activity.

### Spiking neural network architecture and training

To classify mechanical stimulation conditions from model-generated spike trains, we implemented a SNN using the SpikingJelly framework in PyTorch [83]. The SNN received as input the simulated binary spike trains generated by the neuromechanical model and output discrete tactile feature categories corresponding to the stimulation conditions. Each dataset consisted of binary spike trains of dimension *T* × *N* × *K*, as described in the previous section. All samples were normalized in duration by trimming or zero-padding to constant time steps to ensure uniform sequence length. The datasets were then concatenated across conditions and labeled according to stimulation conditions. A 90:10 train-test split was used, with stratified sampling to maintain balanced class distributions. The SNN architecture comprised two fully connected layers with Leaky Integrate-and-Fire (LIF) neurons mediating temporal dynamics (**Fig 5B**). The network structure was as follows:

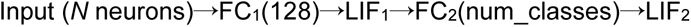

where FC denotes the fully connected layer.

Spike inputs were fed into the network as tensors of shape [*B*, *T*, *N*], where *B* is the batch size. During each forward pass, spikes propagated sequentially through time, and class-wise outputs were accumulated as spike counts over all time steps. The network was trained using the Adam optimizer (learning rate = 10^-3) with cross-entropy loss computed on cumulative output spike counts. Each model was trained for a maximum of 99 epochs with early stopping (training accuracy did not improve for 10 epochs). Training and evaluation were repeated five times per experiment for robustness. The best-performing model checkpoint was automatically saved when the training accuracy improved beyond the set threshold. Final model performance was quantified as the mean and standard deviation of test accuracy across the five training runs. This approach yielded robust estimates of classification reliability under spike variability introduced during stochastic spike generation.

### Insect rearing

Adult *P. americana* were obtained from a laboratory-maintained colony at Penn State University. Cockroaches were reared from eggs to adulthood under controlled conditions: 25 °C, 50% relative humidity, and a 12:12 h light: dark cycle, with continuous access to food and water. Only adult male cockroaches were used for experiments, and all recordings were performed on the same day the animals were collected from the colony.

### Extracellular recordings from antennal nerves

To validate the neural activity predicted by the neuromechanical antenna model, extracellular recordings were performed on the antennal nerves of *P. americana* to obtain ground-truth population responses during controlled antenna-object collisions.

#### Animal preparation

Adult male cockroaches were anesthetized by cooling at 4 °C for approximately 30 minutes prior to surgery. The procedure has been previously described [21]. Briefly, each animal was secured on a custom 3D-printed holder and positioned dorsal side up on a Sylgard gel plate. The thorax and abdomen were fixed using staple pins to minimize body movement while maintaining hemolymph circulation. The head, antennal scape, and pedicel were immobilized using dental silicone (light body dental Alginate Alternative Material; PlastCare USA). The ventral side of the head was oriented parallel to the plate to allow dorsal access to the antenna nerve (**Fig 3A**).

Because the antennal flagellum lacks intrinsic musculature and is innervated solely by sensory afferents [84], the right antenna was positioned horizontally and remained passive during mechanical stimulation. Following tethering, animals were allowed to recover for approximately 20 minutes. To prevent saline leakage, a petroleum jelly barrier was applied around the head, forming a small well that was subsequently filled with cockroach saline [85]. The saline composition was as follows: 9.32 g NaCl, 0.77 g KCl, 0.50 g CaCl_2_, 0.18 g NaHCO_3_, and 0.01 g NaH_2_PO_4_ per liter of distilled water. A small window was cut in the dorsal head cuticle to expose the base of the antennal nerve for electrode placement. The auxiliary heart and antennal vessel were left intact to preserve normal hemolymph flow and physiological conditions during recording.

#### Neural recording module

Following animal preparation, the cockroach was transferred to a Faraday cage to minimize electrical noise during recording. A 32-channel silicon probe (A1×32-Poly3-10mm-25s-177, NeuroNexus, USA) was mounted on a 3-axis micromanipulator and carefully inserted into the exposed antennal nerve at an angle of 70° relative to the horizontal plane. Neural signals were routed through a CM32 adapter connected to an Intan RHD2132 headstage (Intan Technologies, USA) and digitized using an Open Ephys acquisition board (Gen 2) coupled with Open Ephys GUI software for real-time recording and visualization [86]. Neurophysiological data was recorded at a sampling rate of 30,000 Hz.

#### Mechanical stimulation module

Controlled mechanical stimulation of the antenna was achieved using a servo motor-driven 3D-printed arm designed to deliver reproducible collisions. We performed object-antenna collision at three distinct longitudinal regions of antenna: base region (∼10-20% of antennal length), middle region (∼45-55%), and tip region (∼70-80%). Each region was stimulated under two velocity conditions: a slow speed (linear contact velocity ≈ 45 mm/s) and a fast speed (≈ 349 mm/s), which were determined from motor calibration tests and close to the antenna slow exploration speed [43] and cockroach evasive speed [2]. This yielded a total of six stimulation conditions within the same individual: *base-slow*, *base-fast*, *middle-slow*, *middle-fast*, *tip-slow*, and *tip-fast* **(S2 Video)**. For each condition, three repeated trials were conducted to reduce variability and minimize artifacts such as muscle or movement-induced electrical noise. Each trial lasted 8 seconds, followed by at least 30 second inter-trial intervals to minimize sensory adaptation.

#### Synchronization module

To accurately determine the contact onset time between the antenna and the object in each trial, a high-speed camera (Basler acA640-750 um, Basler AG, Germany) was positioned laterally to the animal. The camera sampled at 200 frames per second and transmitted TTL pulses to the Open Ephys acquisition board to embed time-stamped frame markers within the neural data stream. These timestamps enabled precise temporal alignment between mechanical contact events and recorded neural activity during post-processing.

### Post-processing of electrophysiological data

The post-processing pipeline for the extracellular recordings is illustrated in **Fig 3B**. Post-processing was necessary to remove electrical noise originating from antenna muscle activity and to separate overlapping multi-unit signals, ensuring that only well-isolated neuronal units were retained for analysis. Because the recording electrode position remained fixed within each individual across all six stimulation conditions, all trials were concatenated into a single continuous recording using SpikeInterface [87] prior to spike sorting to improve clustering accuracy. Spike sorting was performed using Kilosort 4 [88]. Automated sorting was executed with identical parameters across individuals, using default Kilosort 4 settings except for the following adjustments to match the 32-channel probe configuration: num of channels=32, nblock=0, dminx=21, nearest chans=5, and nearest templates=32. After spike detection and classification, units were categorized based on both automated quality metrics in Kilosort 4 and manual inspection. Units that did not exhibit a clear, physiological waveform shape and a distinct refractory period were discarded. Then the concatenated recording was segmented back into individual trials, and spike timestamps were re-labeled accordingly. Each trial was then temporally aligned to the onset of antenna contact, as identified from the synchronized high-speed camera timestamps.

To quantify neural responses, we converted each unit’s spike train into a continuous firing-rate signal by convolving the binary spike sequence with a Gaussian kernel (σ = 10 ms; kernel width = ±6 σ). The Gaussian-convolved firing rates were then averaged across trials within the same stimulation condition to obtain representative unit activity profiles corresponding to the six defined mechanical stimulations.

## Acknowledgements

We thank Rudolf Schilder and Be Eldash for maintaining the *Periplaneta americana* colony. Research was sponsored by the Army Research Office and was accomplished under Grant W911NF- 23-1-0039 (JMM, KJ) and by the Alfred P. Sloan Research Fellowship FG-2021-16388 (JMM).

## Supporting information

**S1 Fig. Sensitivity analysis of hard-threshold encoding values.** As the encoding threshold increases (red to yellow), the spike density in smaller tactile field regions decreases, leading to a higher compression ratio and a lower recruited ratio. However, higher thresholds significantly delay the time to first-spike. In contrast, the stochastic rate-coding model (black) maintains a superior balance, achieving a high compression ratio (except at threshold = 0.8) and recruitment while consistently delivering faster response times across all simulated conditions. Statistical significance was determined via a Wilcoxon signed rank test comparing the stochastic rate-coding model against each threshold encoding value in each condition. All comparisons for compression and recruited ratios yielded p<0.001.

**S2 Fig. Representative single-unit responses during the base-slow antennal condition.** (A) Gaussian-convolved firing rate of individual units, ordered by the latency to peak firing. Labels represent unit identifiers (e.g. 30-23 denotes the 23rd unit recorded from animal 30) (B) Heatmap of normalized firing rates corresponding to the units shown in (A). (C) Distribution of latencies to maximum firing rate across selected responsive units in base-slow condition.

**S3 Fig. Representative single-unit responses during the base-fast antennal condition.** (A) Gaussian-convolved firing rate of individual units, ordered by the latency to peak firing. Labels represent unit identifiers. (B) Heatmap of normalized firing rates corresponding to the units shown in (A). (C) Distribution of latencies to maximum firing rate across selected responsive units in base-fast condition.

**S4 Fig. Representative single-unit responses during the middle-slow antennal condition.** (A) Gaussian-convolved firing rate of individual units, ordered by the latency to peak firing. Labels represent unit identifiers. (B) Heatmap of normalized firing rates corresponding to the units shown in (A). (C) Distribution of latencies to maximum firing rate across selected responsive units in middle-slow condition.

**S5 Fig. Representative single-unit responses during the middle-fast antennal condition**. (A) Gaussian-convolved firing rate of individual units, ordered by the latency to peak firing. Labels represent unit identifiers. (B) Heatmap of normalized firing rates corresponding to the units shown in (A). (C) Distribution of latencies to maximum firing rate across selected responsive units in middle-fast condition.

**S6 Fig. Representative single-unit responses during the tip-slow antennal condition.** (A) Gaussian-convolved firing rate of individual units, ordered by the latency to peak firing. Labels represent unit identifiers. (B) Heatmap of normalized firing rates corresponding to the units shown in (A). (C) Distribution of latencies to maximum firing rate across selected responsive units in tip-slow condition.

**S7 Fig. Representative single-unit responses during the tip-fast antennal condition.** (A) Gaussian-convolved firing rate of individual units, ordered by the latency to peak firing. Labels represent unit identifiers. (B) Heatmap of normalized firing rates corresponding to the units shown in (A). (C) Distribution of latencies to maximum firing rate across selected responsive units in tip-fast condition.

**S8 Fig. Training performance of SNN using spike trains from the antennal neuromechanical model.** Training and testing accuracy curves for five independent SNN training runs using spike datasets generated with a maximum firing rate of 500 Hz. Solid lines represent training accuracy, dashed lines represent test accuracy, and dotted vertical lines indicate the epoch of optimal performance (maximum train accuracy) for each run.

**S9 Fig. The resting curvature of the biomechanical antenna model was tuned to match the natural antennal curvature measured experimentally (red dots).**

**S10 Fig. Trial-to-trial variability of spike trains generated from identical firing rates.** Spike trains produced by the stochastic spike generator from the same firing rates under the base–slow condition

**S1 Video. Example simulation of the neuromechanical antenna model during antenna–object contact for base-slow condition.**

**S2 Video. Comparison between experimental and simulated antennal dynamics across contact conditions during electrophysiological recordings.**

## Notes

### Competing Interest Statement

The authors declare no competing or financial interests.

### Summary of Updates

The figures were not embedded within the text in the first draft. They are now embedded.

